# Digital spatial profiling of RNA in pancreatic neuroendocrine tumors highlights the distinct profile of α-SMA positive stroma and its impact on surrounding tumor cells

**DOI:** 10.1101/2023.09.05.556304

**Authors:** Helvijs Niedra, Raitis Peculis, Rihards Saksis, Sofija Vilisova, Austra Breiksa, Aija Gerina-Berzina, Julie Earl, Ignacio Ruz-Caracuel, Marta G. Rosas, Vita Rovite

## Abstract

**Objective:** Alpha smooth muscle actin (α-SMA) expression in stroma is linked to the presence of cancer-associated fibroblasts and is known to correlate with worse outcomes in various tumors. In this study, using a digital spatial profiling approach, we characterized the gene expression profiles of the tumor and α-SMA positive stromal cell compartments in pancreatic neuroendocrine tumor (PanNET) tissues.

**Methods:** The profiling was performed in tissues from eight retrospective cases (Three Grade 1, four Grade 2, and one Grade 3) where the segmentation was done based on tissue morphology and synaptophysin (tumor), α-SMA (stroma) marker expression.

**Results:** The stromal cell-associated genes were mainly involved in pathways of extracellular matrix modification, while in tumor cells, the gene expression profiles were primarily associated with the pathways involved in cell proliferation. The comparison of gene expression profiles across all three PanNET grades revealed that heterogeneity is not only present within the tumor but also in the α-SMA positive stromal cells. Furthermore, the comparison of tumor cells adjacent versus non-adjacent to α-SMA positive stromal cells revealed an upregulation of *MMP9* in G3 tumor analysis.

**Conclusions:** Overall, this study provides an in-depth characterization of gene expression profiles in both stroma and tumor cells of PanNETs and outlies potential crosstalk mechanisms.

## Introduction

Neuroendocrine tumors (NETs) are a heterogeneous group of malignancies arising in the neuroendocrine system. The incidence of NETs is rising due to the spread of awareness of the disease and increased accuracy of diagnosis; this trend and estimated prevalence (35 per 100 000) ^1^ highlight the importance of discovering novel biomarkers to improve diagnostic accuracy and optimize disease management.

PanNETs make up about 7% of all NETs and less than 2% of all pancreatic neoplasms ^2^. PanNETs arise from islets of Langerhans, specifically from α- and β-cells of the islets ^3^. PanNETs are classified as non-functioning PanNETs (50% - 90%) and functioning PanNETs (10 %-50%). Non – functioning PanNETs can present high hormone levels but without specific symptoms related to hormone secretion, resulting in nonspecific symptoms such as abdominal pain, weight loss, or mass effect due to tumor or metastasis. Non-functioning, low-grade, and smaller PanNETs usually display a more benign behavior, with slow growth and a better prognosis ^4,5^. The majority (90%) of the PanNETs are regarded as sporadic and arise due to alterations that result in loss function of tumor suppressor genes (*RB1*, *TP53*, *MEN1, TSC2, PTEN, and CDKN2A*) or upregulation of oncogenes (*CCND1, PIK3CA, HRAS, NRAS, and BRAF)* ^6–9^.

Histologically PanNETs are well-differentiated tumors and, based on mitotic count and Ki-67 index, can be further classified into three grades: grade 1 (G1), grade 2 (G3), and grade 3 (G3) ^10^. G1 and G2 tumors usually present as small (<3 cm) hypervascular masses, while G3 tumors typically appear as heterogeneous masses larger than 3 cm with areas of necrosis and lesser vascular density than lower-grade tumors ^10^. Grading classification based on the Ki-67 index seems to be not sufficient. Consequently, the inclusion of mutation analyses is proposed for the upcoming classifications ^5^.

One of the major components of any tumor is its microenvironment, also known as the tumor microenvironment (TME). It is a highly heterogeneous environment that consists of four major components: immune cells, stromal cells, blood vessels, and extracellular matrix ^11^. Alongside the other components in TME, an important role is played by stroma, which mainly consists of fibroblasts, endothelial cells, and extracellular matrix ^12^. The stromal cells can either hinder tumor growth or promote it by upregulating angiogenesis, extracellular matrix remodeling, immune suppression, and altering the metabolism of the cells within TME ^12^. Recent studies have shown that the upregulation of alpha-smooth muscle actin (α-SMA) in tumor stromal compartment correlates with worse outcomes and therapeutic resistance in various tumors ^13,14^. This could be attributed to the fact that α-SMA is highly expressed by transformed fibroblasts known as cancer-associated fibroblasts (CAFs) ^15^, which can harbor significant tumor-promoting functions ^16^. The molecular aspects of stroma have also been studied within NETs, where it has been shown that NET cells can recruit stromal fibroblasts by transforming them into α-SMA expressing CAFs via the secretion of TGF-β and platelet-derived growth (PDGF) factors ^17^.

Due to the rarity of PanNETs, there are only a few studies that have characterized the overall transcriptomic landscape of PanNETs ^18,19^, and despite the evidence of existing crosstalk between stromal cells and NET cells, the underlying molecular mechanisms on the transcriptomic level remain largely understudied ^17^. More so, the currently existing transcriptomic studies are limited by the bulk RNA sequencing approach, where the spatial and cellular composition information is lost ^20^. Therefore, within this study, we employed GeoMx digital spatial profiling (DSP). A technology that allows us to extract the gene expression profiles from specific tissue compartments, which can be identified using morphology markers ^21^. As a result, we characterized the gene expression profiles of 1800 cancer-associated genes in PanNET cells surrounding α-SMA positive (α-SMA+) stromal cells and islet/acinar compartments of the adjacent normal tissue. Furthermore, we also evaluated the differences in gene expression profiles between tumor cells that are adjacent to α-SMA+ stromal cells and tumor cells that are non-adjacent (distant) from α-SMA+ stromal cells.

## Materials and Methods

### Study cohort

A retrospective series (2017–2021) of eight surgically resected non-functioning primary sporadic PanNETs from two sites (Latvia and Spain) were investigated. The cohort included five males and three females with a median age at surgery of 58 years (range: 36–77 years). Written consent was obtained from each patient individually, and the study and inclusion of patients’ samples were approved by: The Central Medical Ethics Committee of Latvia (protocol No. 1.1–2/67) and Ramon y Cajal Ethical and Scientific Committees (protocol No. 196-19), samples were provided by the BioBank Hospital Ramón y Cajal-IRYCIS (National Registry of Biobanks B.0000678), integrated into the Biobanks and Biomodels Platform of the ISCIII (PT20/00045). All patients’ tumor samples were synaptophysin positive, and none of the patients had received medical treatment before the surgery. Additional collected data included the sex of the patients, grade and differentiation of the tumor, Ki-67 index, stage of the disease, and patient age at surgery (Table 1). All cases were classified by grade of the tumor according to WHO 2019 criteria [doi: 10.1111/his.13975]. In total, three G1, four G2 patients, and one G3 patient were recruited, and FFPE samples of the tumor were obtained for DSP analysis. Six out of eight patients also had available FFPE samples of adjacent normal pancreatic tissues (Table 1).

**Table 1.**
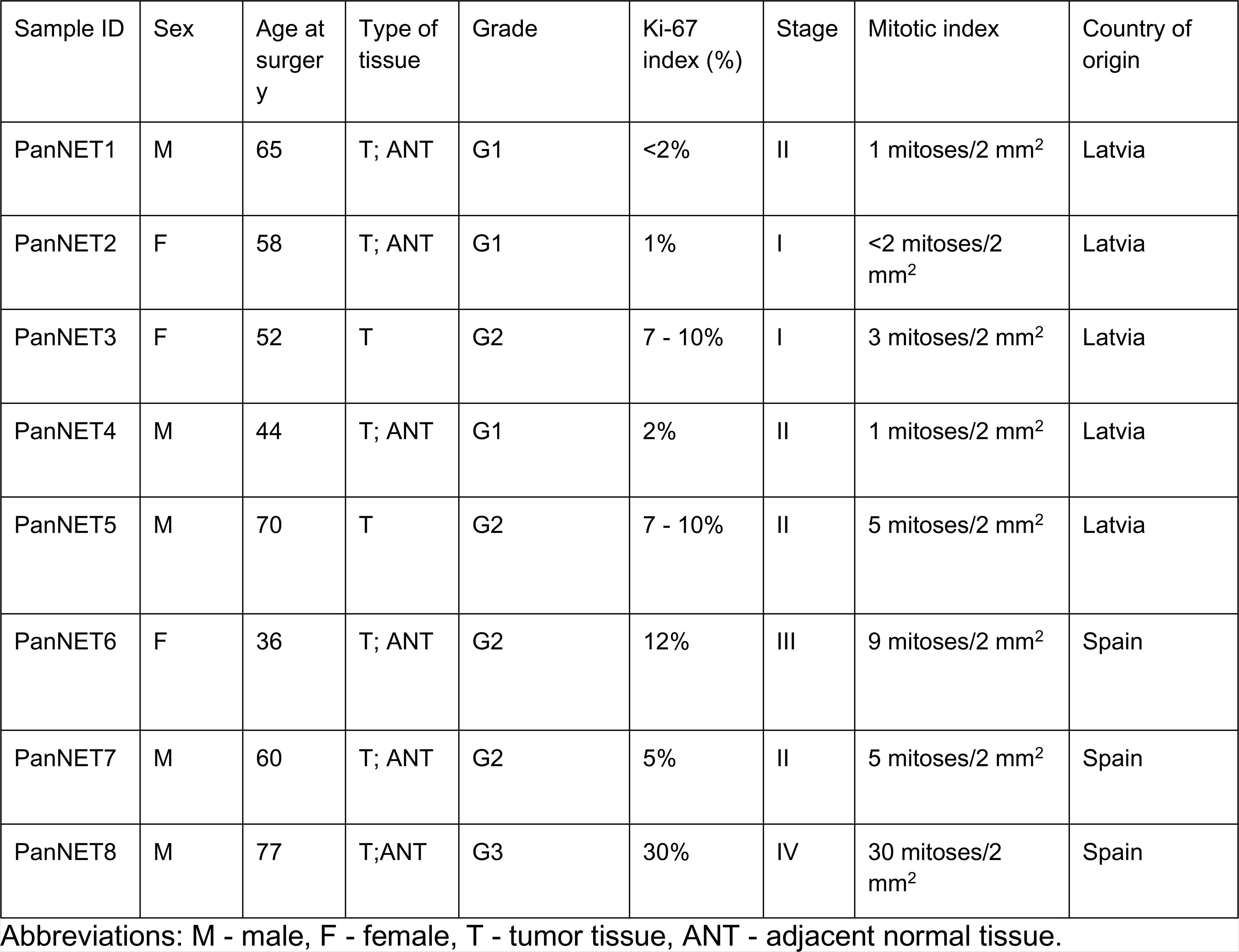
Clinical characteristics of the patients recruited in the study.

### Slide preparation for DSP analysis

The slides were incubated for 3 hours at 65oC for paraffin removal and subsequently loaded onto a Leica BOND RX for tissue rehydration, heat-induced epitope retrieval (ER2 for 20 minutes at 100°C), and proteinase K treatment (0.1 μg/ml for 15 minutes at 37°C). The tissue sections were then hybridized with the Cancer Transcriptome Atlas (CTA) probes overnight at 37°C. Following 2x 5 min stringent washes (1:1 4x SSC buffer & formamide), the slides were blocked and then incubated with morphology marker antibodies to guide region of interest (ROI) selection: α-SMA (488 channel, 53-9760-82, ThermoFisher) and Synaptophysin (647 channel, ab196166, Abcam). Syto83 (532 channel, S11364, Invitrogen) was used as a nuclear stain after region of interest (ROI) selection. For segmentation, UV light was directed by the GeoMx at each area of illumination (AOI), releasing the RNA ID and UMI-containing oligonucleotide tags from the CTA probes for collection and preparation for sequencing. Illumina i5 and i7 dual indexing primers were added to the oligonucleotide tags during PCR to index each AOI uniquely. AMPure XP beads (Beckman Coulter) were used for PCR purification. Library concentration was measured using a Qubit fluorometer (Thermo Fisher Scientific), and quality was assessed using a Bioanalyzer (Agilent). Sequencing was performed on an Illumina NovaSeq 6000. The stained sections were analyzed, and ROIs for sequencing analysis were selected by the GeoMx and a pathologist specialized in neuroendocrine tumors.

### Processing and analysis of DSP data

The sequencing FASTQ files were processed using the GeoMx® NGS Pipeline to acquire GeoMX DSP Analysis Suite compatible DCC files. Following this, the DCC files were analyzed on GeoMX DSP Analysis Suite, which included data quality control, filtering, normalization, and differential expression analysis. For the data quality control step, the reads from all 105 AOIs were inspected for raw reads aligned (cutoff set at 80%), sequencing saturation (cutoff set at 50%), negative probe count geomean (cutoff set at five counts), and minimum nuclei count (cutoff set at 200 nuclei). As a result, 78 AOIs were flagged for negative probe count geomean below 5, six were flagged for aligned read percentage below 70%, and 30 AOIs were flagged for nuclei count below 200. Probe QC was also performed, and any probes that did not meet the following criteria: (geomean probe in all AOIs) / (geomean probes within target) ≤ 0.1, and failed Grubbs’ outlier test in ≥20% of AOIs, were removed from target count calculation. Following this, we also performed AOI and target filtering steps. For AOI filtering, any AOIs that had less than 5% of targets above default (higher of LOQ and user-defined value of 2) expression threshold were filtered. The filtering was also applied to targets, and any targets that were present in less than 3% of AOIs, were removed from further analysis. After the quality control and filtering steps, 466 genes and one AOI were excluded from further analysis leaving a final count file containing information on 1482 genes and 104 AOIs which was used for the downstream analysis (Supplementary Table 12). Following this, the raw count data were normalized using the Q3 normalization method (Supplementary Table 13). The metadata for each AOI can be found in Supplementary Table 1. Dimension reduction analysis was performed using the GeoMx DSP analysis suite plugin - DimensionReduction (v1.1). Spatial deconvolution analysis was performed in R (v4.2.2) using a Bioconductor package SpatialDecon (v1.8.0) with the inbuilt SafeTME cell profile matrix ^68^. To evaluate the changes in expression between different AOI types, a linear mixed model analysis was performed in GeoMx DSP Analysis Suite, and volcano plots depicting the results were made using the GeoMx Analysis Suite plugin - Volcano Plot (v1.2).

### Pathway enrichment and protein-protein interactions analyses

Differentially expressed genes (DEGs) with Log2FC > 0.5 and FDR < 0.05 values were subjected to pathway enrichment analysis and protein-protein interaction (PPI) analysis using STRING database (v11.5) ^69^ and stringApp plugin (v2.0.1) in Cytoscape (v3.9.1). Dot plots depicting the enriched pathways were made using R (v4.2.2) package ggplot2 (v3.4.0), and PPI networks were constructed in Cytoscape using additional plugins: NetworkAnalyzer (v4.4.8), LegendCreator (1.1.6) and enhanced graphics (1.5.5)

## Results

### Overview of AOIs analyzed by the digital spatial profiler

Based on immunofluorescence and morphology, we selected 45 regions of interest (ROIs) within eight tumor tissue samples based on the proximity of the tumor cells to the α-SMA+ stroma (detailed information on each tumor sample can be found in Table 1). As a result, 24 ROIs contained tumor cells adjacent to α-SMA+ stromal cells, and 21 ROIs contained tumor cells without adjacent α-SMA+ stromal cells. The tumor ROIs adjacent to α-SMA+ stroma were segmented into two AOIs based on immunofluorescence signals. This yielded 24 AOIs containing tumor cells and 23 AOIs containing α-SMA+ stromal cells (one AOI was dropped due to a low number of targets above the defined expression threshold) (Figure 1 A). This amounted to a total of 68 AOIs from 45 ROIs within tumor tissue samples. Using DSP, we also investigated normal pancreatic tissue and selected 18 ROIs containing Islets of Langerhans and 18 ROIs containing acinar compartment, which amounted to a total of 36 AOIs from 36 ROIs. As a result, the gene expression was assessed in AOIs; more detailed information on the AOIs analyzed can be found in Supplementary Table 1. By applying the T-distributed neighbor embedding (t-SNE) dimensionality reduction, we can observe segregation by AOI type as a distinction between tumor, α-SMA+ stroma, and normal cell AOIs can be observed (Figure 1 B). Following this, the gene expression data of all 68 tumor AOIs were subjected to cell type deconvolution analysis to estimate the relative abundance of cell types within each AOI. By examining the proportion of fitted cells (Figure 1 C), it can be observed that α-SMA+ stroma AOIs have a higher proportion of stromal cells than tumor AOIs. As expected, within the tumor specific AOIs, the most abundant cell type was PanNET cells. We also observed the second most abundant cell type within tumor tissue was T cells. A complete list of cell type counts and proportions in tumor tissues can be found in (Supplementary Table 2)

**Figure 1.**
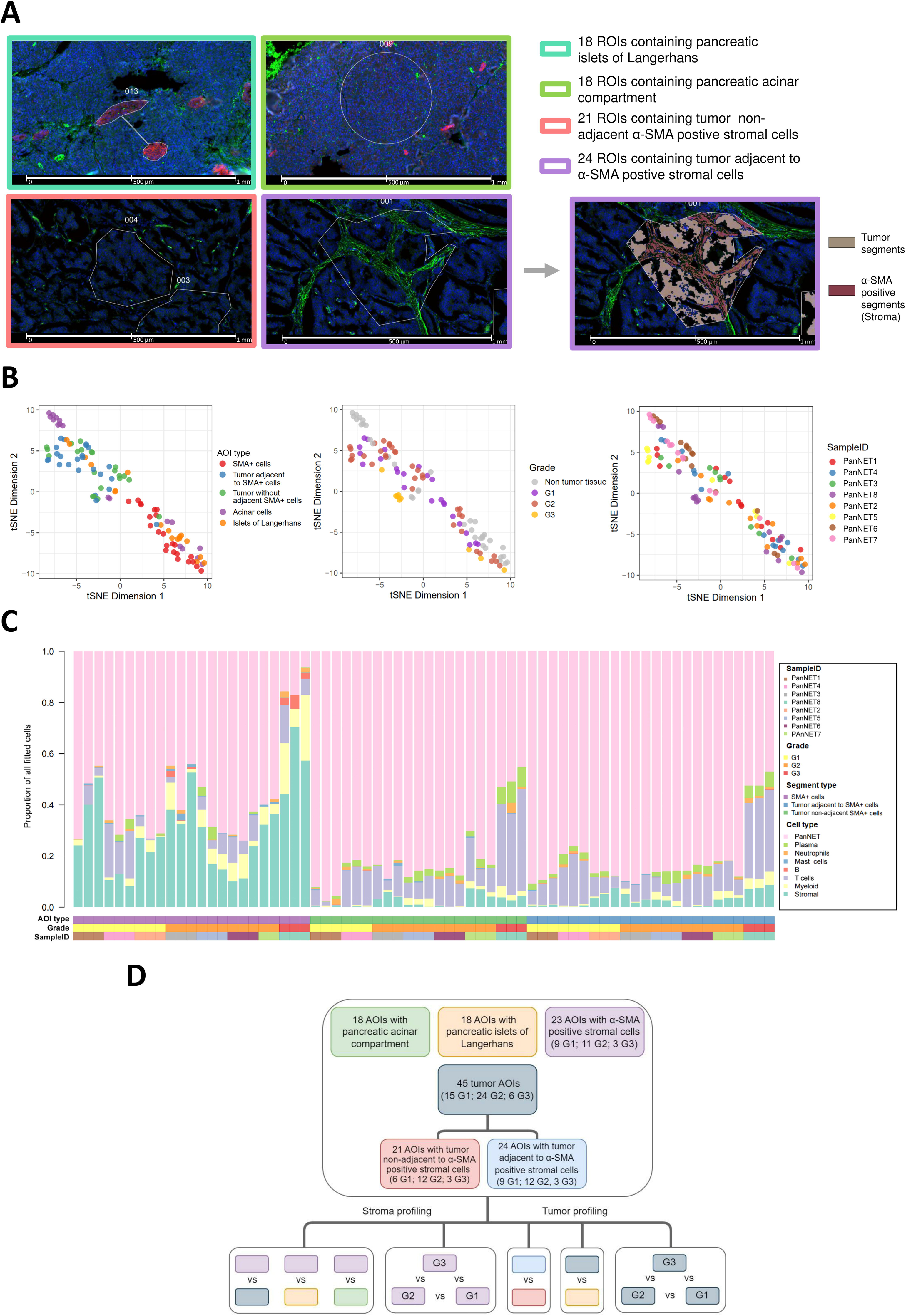
Study design for digital spatial profiling of three G1, four G2, and one G3 pancreatic neuroendocrine tumor tissue samples. **A** – represents the strategy that was used to select regions of interest (ROI) and areas of illumination (AOI). ROIs containing islets of Langerhans (18x) and acinar compartment (18x) within adjacent normal tissues were selected based on the presence of synaptophysin and morphology, no further segmentation was performed, and AOI count was the same as the ROI count (36x). Within the tumor, ROI selection was based on the proximity of alpha-smooth muscle actin-expressing stromal cells (α-SMA+ stroma). As a result, 21 ROIs with tumor non-adjacent to α-SMA+ stroma were selected, and no further segmentation was performed here, yielding 21 AOIs. Following this, 24 ROIs adjacent to α-SMA+ stroma were selected, and segmentation was performed based on the α-SMA signal yielding 24 tumor AOIs and 24 α-SMA+ stroma AOIs. **B** - T-distributed neighbor embedding (t-SNE) dimensionality reduction results with 104 sequenced AOIs. **C** - Stacked bar chart from spatial deconvolution analysis displaying the proportion of all fitted cells in each AOI from tumor tissue. **D** - study design of the subsequent differential expression analyses with 104 AOIs. For tumor and α-SMA+ stromal AOIs, the comparisons were done for each tumor grade separately.

### Identification of genes associated with SMA+ stroma

To identify genes specific to α-SMA+ stroma within PanNET tissues, we carried out differential expression analysis where α-SMA+ stroma AOIs of each tumor grade were compared to tumor AOIs and acinar compartment/islet cell AOIs from normal tissue. After the differential expression analyses, we identified the sets of overlapping genes between these comparisons to further increase the odds of discovering the list of genes that are specifically associated with α-SMA+ stroma (Supplementary Table 3). As a result, in the comparisons of α-SMA+ AOIs from G1 tumor (Figure 2A), we identified a set of 54 overlapping genes (Figure 2B), of which 42 had a positive, seven had a negative, and three had a mixed fold change direction in all three comparisons. Further subjecting the set of 54 overlapping genes to STRING functional enrichment analysis showed 32 overrepresented Reactome pathways (Supplementary Table 4). The top 15 pathways are listed in Figure 2C. To further expand on the interactions between overlapping genes, a STRING network of physical protein associations was constructed, which revealed 47 interactions between 34 proteins with protein-protein interaction (PPI) enrichment p-value of 1.0 × 10^−16^. In the G2 scenario (Figure 3 A), 105 overlapping genes were found, which was substantially higher than in the G1 scenario (Figure 3B). Similarly to the G1 scenario, here we also observed a uniform fold change direction of the overlapping genes as 53 genes showed positive, 41 negative, and 11 mixed fold change directions in all three comparisons. Here the STRING enrichment analysis revealed 190 overrepresented Reactome pathways (Supplementary Table 4). The top 15 pathways are listed in Figure 3 C. In the network of physical protein associations, 89 interactions were found between 54 proteins (Figure 3 D) with a p-value of 6.622 × 10^−10^, indicating that proteins of overlapping genes function as a biological group. Lastly, within the G3 comparison scenario (Figure 4 A), the amount of overlapping genes (Figure 4 B) was the highest (121 genes). There was also a uniform fold change direction across all three comparisons as 71 genes showed positive, 35 negative, and 15 mixed fold change directions in all three comparisons. Amongst the list of 121 overlapping genes, 90 overrepresented Reactome pathways were discovered (Supplementary Table 4), and the top 15 pathways are listed in Figure 4 C. Within the network of physical protein associations, the interactions were more pronounced compared to the G2 overlapping gene network as 137 interactions were observed between 77 proteins with a p-value of 1 × 10^−16^.

**Figure 2.**
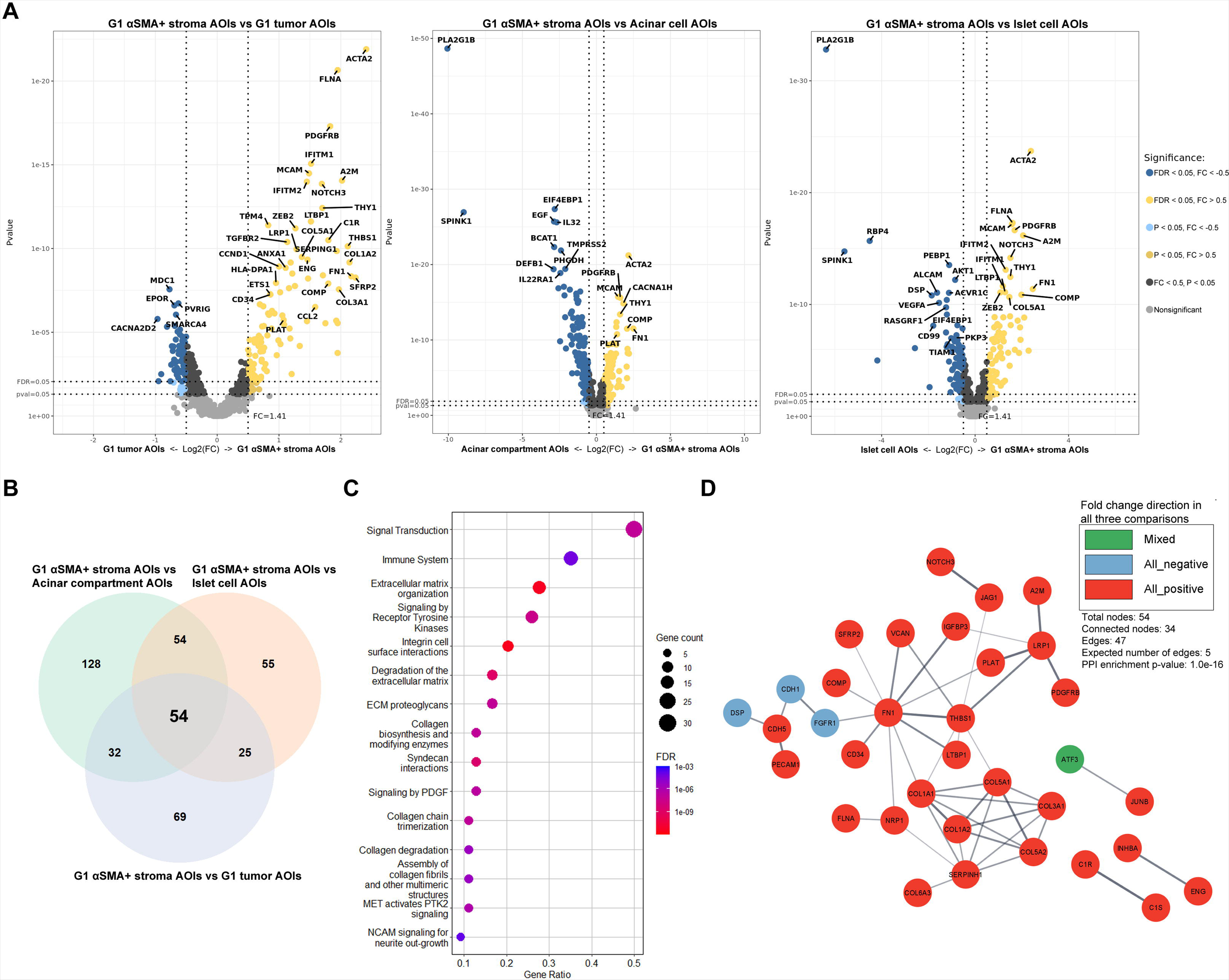
Analysis of alpha-smooth muscle actin positive (α-SMA+) stroma AOIs from G1 tumor tissues. **A** - volcano plots depicting the Log_2_ fold changes and p-values from differential expression analyses where a set of nine G1 α-SMA+ stroma AOIs were compared against sets of 15 G1 tumor, 18 acinar compartment, and 18 islet cell AOIs. **B -** Venn diagram depicting a set of 54 overlapping differentially expressed genes between all three comparisons, which were used as an input for pathway enrichment analysis. **C -** Dot plot depicting the top 15 overrepresented Reactome pathways according to FDR value sorted by the gene count related to the term. **D -** STRING network of physical protein associations showing physical interactions between proteins of differentially expressed genes.

**Figure 3.**
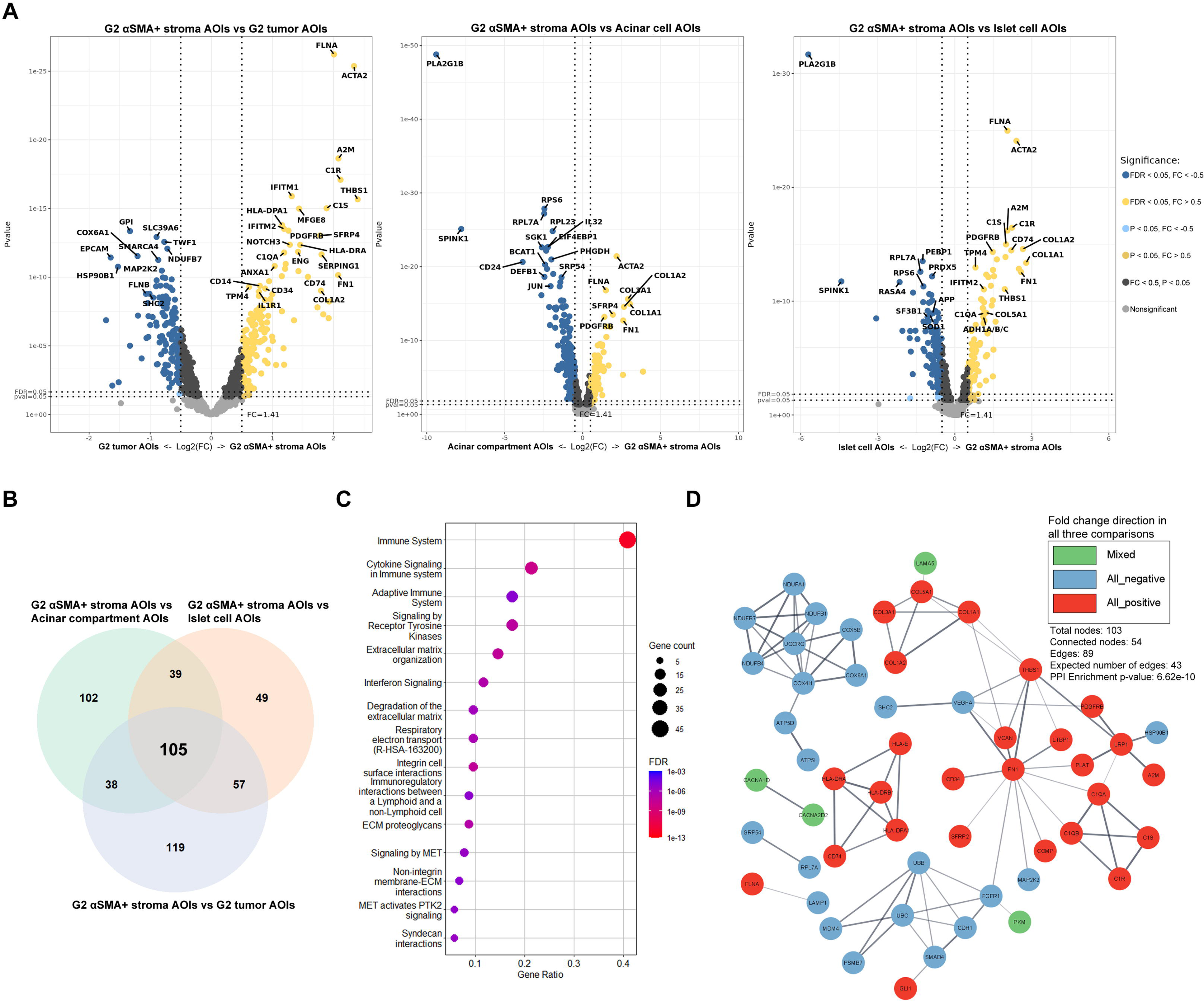
Analysis of alpha-smooth muscle actin positive (α-SMA+) stroma AOIs from G2 tumor tissues. **A** - volcano plots depicting the Log_2_ fold changes and p-values from differential expression analyses where a set of 11 G2 α-SMA+ stroma AOIs were compared against sets of 24 G2 tumor, 18 acinar compartment, and 18 islet cell AOIs. **B -** Venn diagram depicting a set of 54 overlapping differentially expressed genes between all three comparisons, which were used as an input for pathway enrichment analysis. **C -** Dot plot depicting the top 15 overrepresented Reactome pathways according to FDR value sorted by the gene count related to the term. **D -** STRING network of physical protein associations showing physical interactions between proteins of differentially expressed genes.

**Figure 4.**
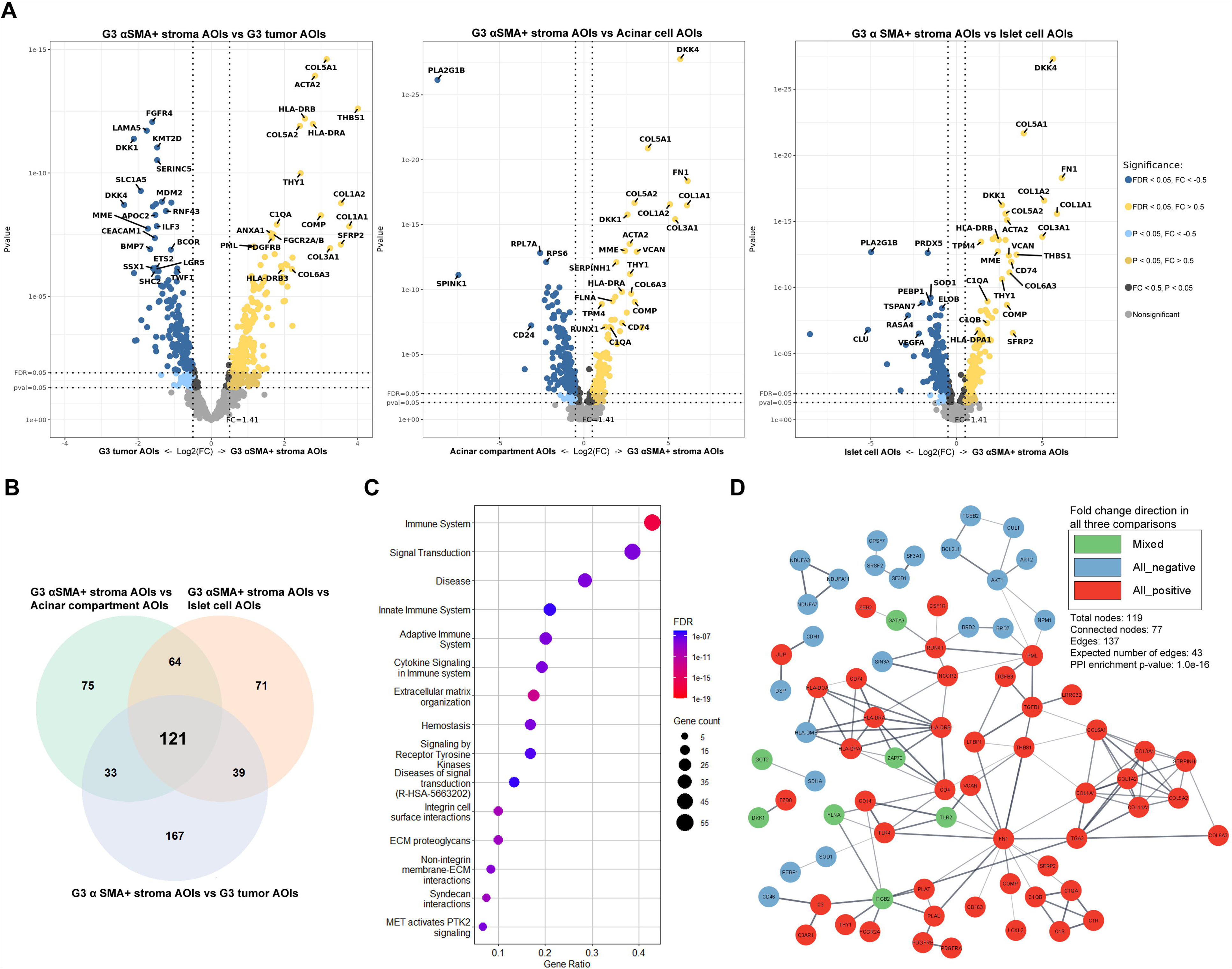
Analysis of alpha-smooth muscle actin positive (α-SMA+) stroma AOIs from G3 tumor tissues. **A** - volcano plots depicting the Log_2_ fold changes and p-values from differential expression analyses where a set of three G3 α-SMA+ stroma AOIs were compared against sets of six G3 tumor, 18 acinar compartments, and 18 islet cell AOIs. **B -** Venn diagram depicting a set of 54 overlapping differentially expressed genes between all three comparisons, which were used as an input for pathway enrichment analysis. **C -** Dot plot depicting the top 15 overrepresented Reactome pathways according to FDR value sorted by the gene count related to the term. **D -** STRING network of physical protein associations showing physical interactions between proteins of differentially expressed genes.

In all three scenarios, the top 15 pathways associated with the overlapping DEGs were mainly related to the events of the immune system, extracellular matrix organization, and MET signaling (Figures 2C, 3C, 4C). Upon further inspection of STRING protein networks, we observed that in G2 and G3 tumors, the immune system pathways were especially pronounced as a cluster of tightly interacting genes related immune system were found: *CD74, HLA-E, HLA-DRA, HLA-DPA1, HLA-DRB1* in G2 (Figure 3D) and *CD74, HLA-DRA, HLA-DPA1, HLA-DRB1, HLA-DOA, HLA-DMB, ZAP70* in G3 (Figure 4D). Following this, we also identified clusters containing a multitude of tightly interacting collagen genes which were upregulated in α-SMA+ stroma across all three grades: *COL1A1, COL5A1, COL1A2, COL3A1, COL5A2* in G1 (Figure 2D), *COL1A1, COL5A1, COL1A2, COL3A1* in G2 (Figure 3D), *COL1A1, COL5A1, COL1A2, COL3A1, COL5A2, COL6A3, COL11A* in G3 (Figure 4D). The pathway enrichment analysis shows that these genes are exactly involved in pathways regarding extracellular matrix organization and MET signaling events. In combination with the aforementioned immune-related genes, collagen genes were also involved in immune system pathways. It can also be observed that all the collagen proteins are strongly linked to the *SERPINH1* gene (Figure 2D, Figure 4D), which according to the STRING database, is a serine protease inhibitor and plays a role in collagen processing by acting as a chaperone.

Lastly, we also looked at the expression of various ligands secreted by stromal cells, particularly the CAF component of stroma, which has been listed in multiple secretome studies and reviews ^22–28^. Here we also focused on reporting only those factors which were differentially expressed (FDR < 0.05) in all three comparisons. Across all three tumor grades, we observed upregulation of two factors: *TGFB1* (TGF-β) and *FN1* (fibronectin) in α-SMA+ stroma AOIs when compared to all other tissue type AOIs (Figure 5 A, B, C). Additionally, in the G1 and G3 groups, one additional factor was upregulated: *IGFBP3* (IGF-Binding Protein 3) in the G1 group and *FGF8* (fibroblast growth factor 8) in the G3 group. In the G2 group, two additional factors were differentially expressed where *CCL5* (RANTES) was upregulated, and *VEGFA* (vascular endothelial growth factor) was downregulated. Furthermore, some studies have shown that *FN1* and *CCL5* ^29–31^ gene expression positively correlates with immune cell infiltration within the tumor. Therefore, we performed a linear regression and Pearson correlation analysis to investigate this relationship in our samples with *FN1* expression data and cell count data acquired from deconvolution analysis. As a result, we observed that *FN1* expression in α-SMA+ stroma had a statistically significant positive correlation with both myeloid cell and T cell levels in tumor AOIs adjacent to α-SMA+ stroma (Figure 5 D). We also evaluated the correlation in stromal AOIs themselves, but here a statistically significant correlation was observed only between *FN1* and myeloid cell levels (Figure 5E). As for the *CCL5* gene, although positive correlations were observed in all scenarios, none of them were statistically significant.

**Figure 5.**
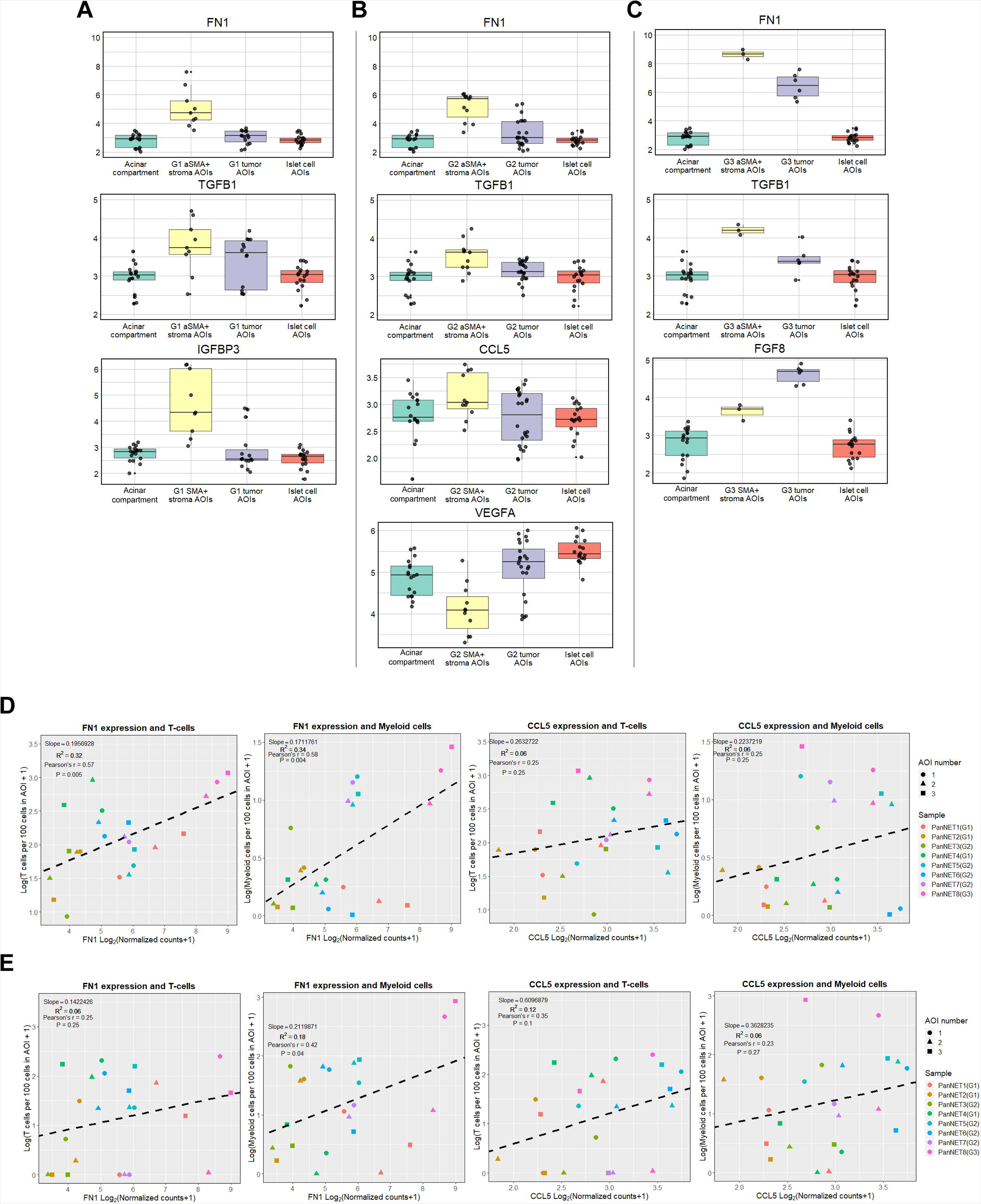
**A, B, C** - box plot visualization of overlapping differentially expressed genes related to proteins secreted by cancer-associated fibroblasts. The box plots represent the genes that were differentially expressed in all three comparisons: α-SMA+ stroma versus 1) tumor, 2) acinar compartment, and 3) islet cell comparisons. **A -** grade 1 tumors, **B -** grade 2 tumors, **C -** grade 3 tumors. Y axis - Log_2_(Normalized counts + 1). **D, E** - Scatter plots with linear regression trendlines representing a correlation between FN1/CCL5 expression and T-cell/myeloid cell levels in **D -** tumor AOIs adjacent to α-SMA+ stromal cells, **E -** α-SMA+ stromal cells AOIs. The X-axis represents Log_2_(Normalized counts + 1) of the target gene, and the Y-axis represents Log(Cell counts per 100 cells in AOI +1) of the target cell type.

### Comparison of α-SMA+ stroma segments across tumor grades

Our next goal was to understand the differences in gene expression profiles of α-SMA+ AOIs between different tumor grades. As a result, we carried out differential expression analysis where α-SMA+ AOIs were compared against each other depending on the tumor grade (G1, G2, G3) (Supplementary Table 5). When G2 α-SMA+ AOIs were compared to G1, a total of 58 DEGs (22 upregulated, 35 downregulated) were identified at FDR < 0.05 and Log2FC > 0.5 cutoffs. In STRING enrichment analysis, 11 Reatcome pathways were associated with these 58 DEGs (Figure 6 A, Supplementary Table 6). Amongst these results, we found three notable terms belonging to NOTCH signaling. The genes in association with the NOTCH signaling pathways were *APH1B, JAG1, and UBC,* which were all upregulated in G1 tumors compared to G2. The STRING network of physical protein associations showed a relatively low amount of associations between the proteins of these 58 DEGs, as the network consisted of 16 interactions between 23 nodes with a PPI enrichment p-value of 0.41. When we compared the G3 tumor to both G1 and G2, a total of 109 (66 upregulated, 43 downregulated) (Figure 6 B) and 63 (45 upregulated, 18 downregulated) (Figure 6 C) DEGs were found. Since G3 and G1 tumors are histologically furthest apart, the highest amount of DEGs was expected in this comparison. In G3 vs. G1 comparison, DEGs subjected to STRING enrichment analysis showed an association with 129 pathways, while in the G3 vs. G2 list of DEGs, 39 pathways were overrepresented (Supplementary Table 6). In both comparisons, the top 15 enriched pathways mainly belonged to events of extracellular matrix organization, immune system, and MET signaling. Also, in both G3 comparisons, statistically significant networks of STRING physical protein interactions were discovered. In G3 vs. G1, the network consisted of 61 proteins with 98 interactions with a PPI enrichment p-value of 3.3 × 10^−10^, and in G3 vs. G2, the network consisted of 35 proteins with 51 interactions with a PPI enrichment p-value of 1 × 10^−16^. As depicted by the Venn diagram (Figure 6 D), it can be concluded that G3 tumor stroma is more different from G1 tumor stroma than from G2, showing that the PanNET stroma also changes as the tumor progresses through grades. Further looking at the CAF-associated markers presented in Figure 5 (*FN1* and *TGFB1*), we observed an increase of FN1 expression in G3 tumor when compared to G2 or G1 counterparts. As for the *TGFB1,* the expression of this gene in stroma did not vary significantly between tumor grades.

**Figure 6.**
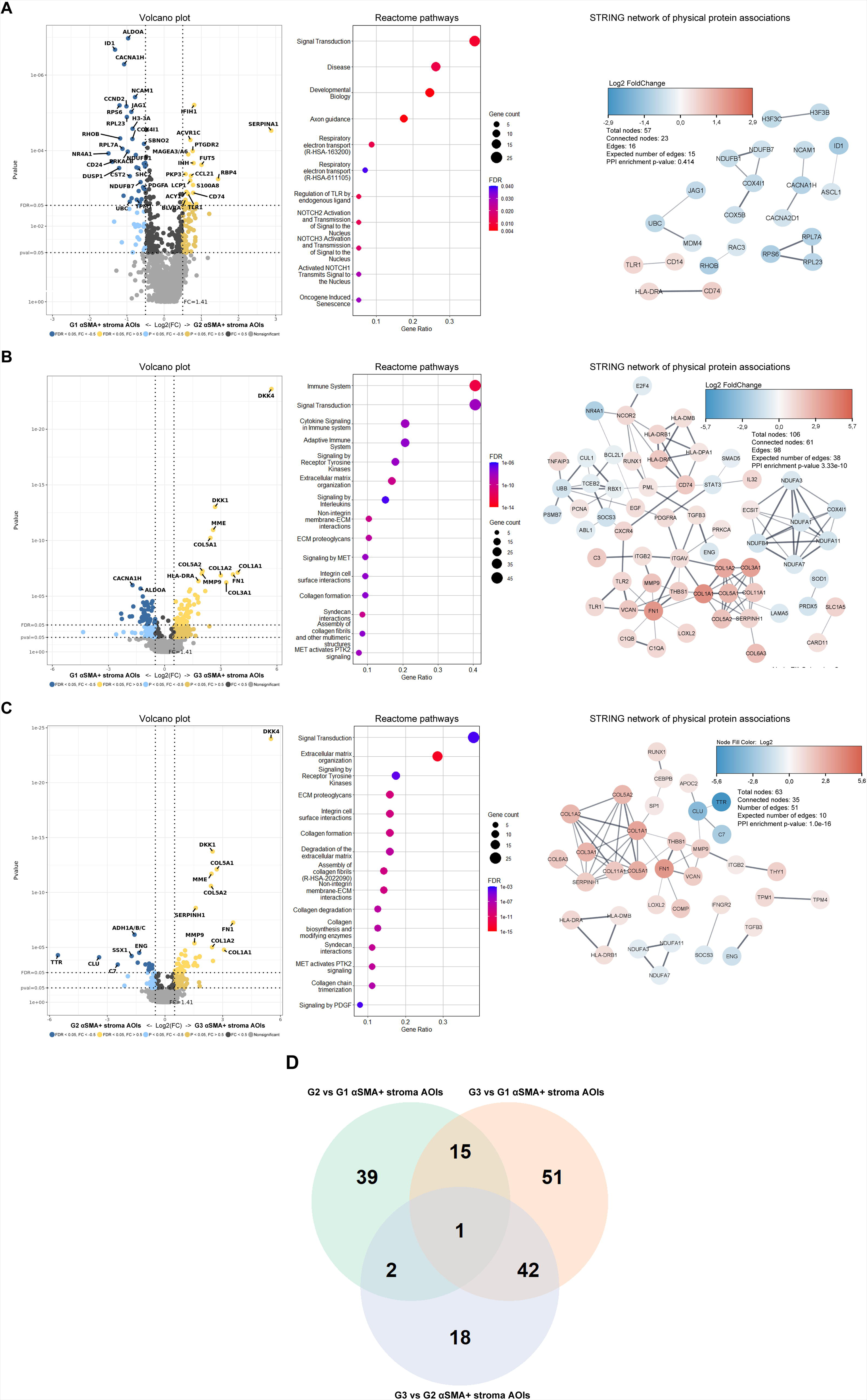
Comparison of alpha-smooth muscle actin positive (α-SMA+) AOIs between different grades of the tumor. **A - C** figures contain volcano plots depicting the Log_2_ fold changes and p-values of the assessed genes, dot plots of the top 15 Reactome pathways according to FDR value which is sorted by gene count related to the term, and lastly, STRING network of physical protein associations depicting the physical interactions between proteins of DEGs. **A** - G2. vs G1; **B** - G3 vs. G1. In this comparison, four out of 109 input genes were excluded from the STRING enrichment analysis due to missing information in the STRING database; **C** G3 vs. G2. **D** - Venn diagram depicting the overlapping DEGs between three comparisons.

### Changes in tumor gene expression in relation to the proximity of α-SMA+ stromal cells

To assess whether there are differences in tumor cells in regards to their proximity to the α-SMA+ stromal cells, the tumor AOIs from ROIs that encompassed both tumor and α-SMA+ stromal cells were compared against tumor AOIs from ROIs that were distant from any α-SMA+ stroma cells (Supplementary table 7). Across all three grades, the differences in gene expression were marginal (Figure 7). At the FDR cutoff at 0.05 and Log_2_FC cut off at 0.5, only one DEG was found (*MMP9*) in a G3 tumor which was upregulated in tumor AOIs adjacent to the α-SMA+ stroma. By taking into account only the unadjusted p-values and without Log_2_FC cutoff in G1 tumors, changes were observed for 81 genes (55 upregulated, 26 downregulated). In G2 tumors, the changes were far less pronounced as only for nine genes a change in expression was observed. Lastly, in G3 tumor, aside from *MMP9*, 15 additional genes (5 upregulated, ten downregulated) had a p-value of < 0.05.

**Figure 7.**
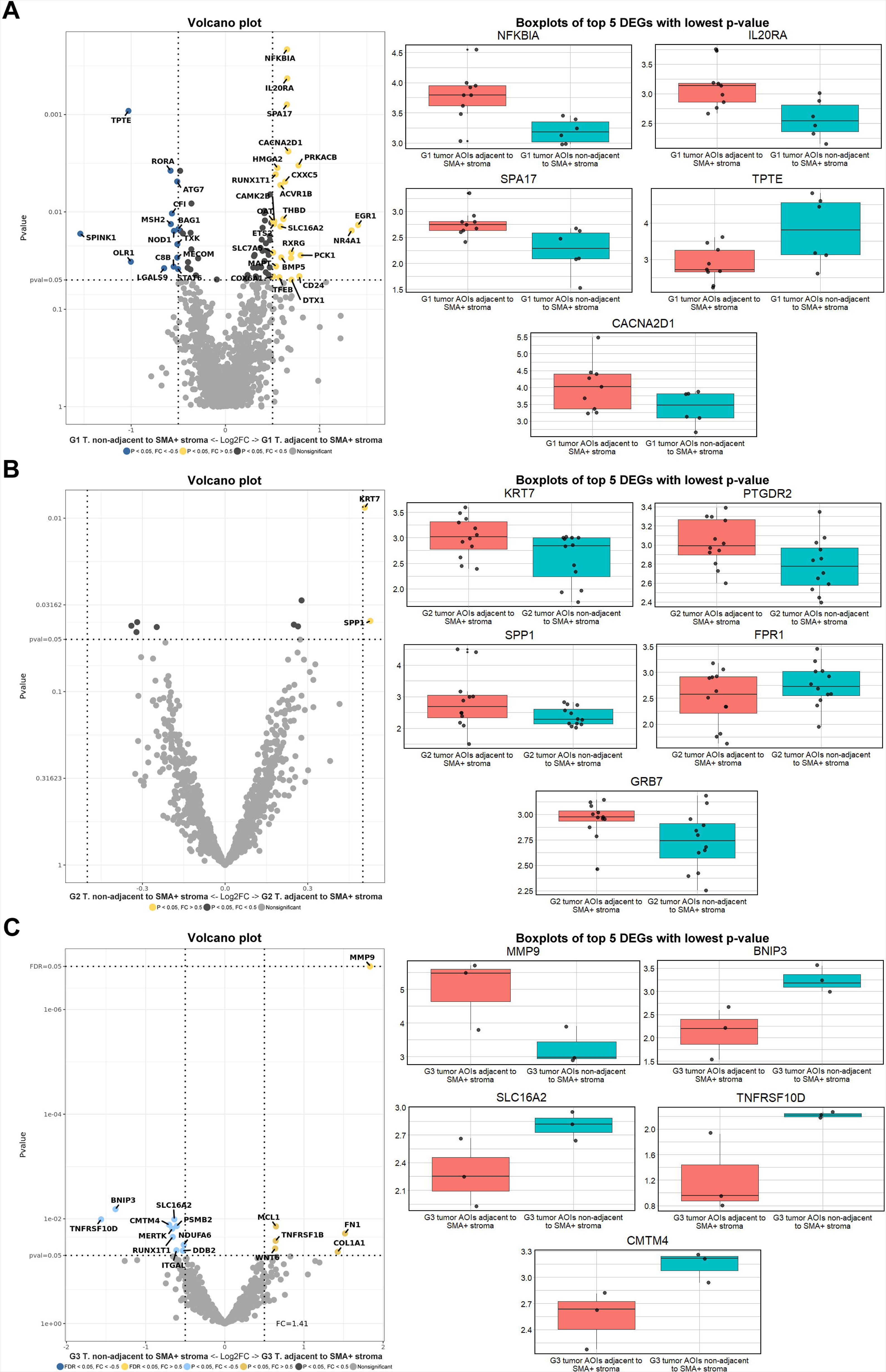
Results of differential expression analysis between tumor adjacent to alpha-smooth muscle actin positive (α-SMA+) stroma vs. tumor non-adjacent to α-SMA+ stroma across all three tumor grades. The results contain volcano plots depicting the Log_2_ fold changes and p-values of assessed genes. The boxplots on the right section depict the Log_2_(Normalized counts + 1) transformed normalized count data of five differentially expressed genes with the lowest p-value. **A -** G1 tumors comparison, **B** - G2 tumors comparison, **C -** G3 tumor comparison.

### Identification of altered genes in PanNET cells in comparison to pancreatic islet cells

To identify genes that are specifically associated with PanNET cells, we compared the AOIs containing PanNET cells against AOIs containing pancreatic islets of Langerhans (Supplementary Table 8). Since PanNETs arise from the islet cells, this allows us to more precisely assess the changes that are related to tumorigenesis rather than to the difference in cell types. In the list of DEGs with FDR < 0.05 and Log_2_FC > 0.5 (volcano plots in Figure 8 A, B, C), the first notable finding here is that amongst all three grades, we can observe 26 overlapping DEGs (Figure 8 D) of which five genes (*RBP4*, *SPINK1, PLA2G1B, EGR1, and CD99*) showed a consistent downregulation (Figure 8 E). Additionally these were also amongst the top 10 downregulated genes in all three comparisons. By looking at each comparison individually at FDR < 0.05 and Log2FC > 0.5 thresholds in G1 tumor AOIs comparison against islet cell AOIs, we identified 114 differentially expressed genes (89 downregulated, 25 upregulated) which showed statistically significant associations with 51 Reactome pathways in STRING enrichment analysis (Supplementary table 9). Amongst the top 15 pathways (Figure 8 A), we found RAF/MAP kinase cascade, MAPK signaling cascades, and PIP3 activates AKT signaling, which are all linked to cell proliferation and survival. Another unique pathway was signaling by VEGF, which is involved in angiogenesis and has been extensively studied in neuroendocrine tumors ^32^. The majority of DEGs related to this pathway (*ITGAV, KDR, VEGFB, AKT1, VEGFA*) were downregulated in tumor cells except for *HSBP1* and *SHC2*. Further analyzing physical interactions between DEGs, a STRING network of physical protein associations showed 75 interactions between 64 proteins with an enrichment p-value of 4.88 × 10^−9^.

**Figure 8.**
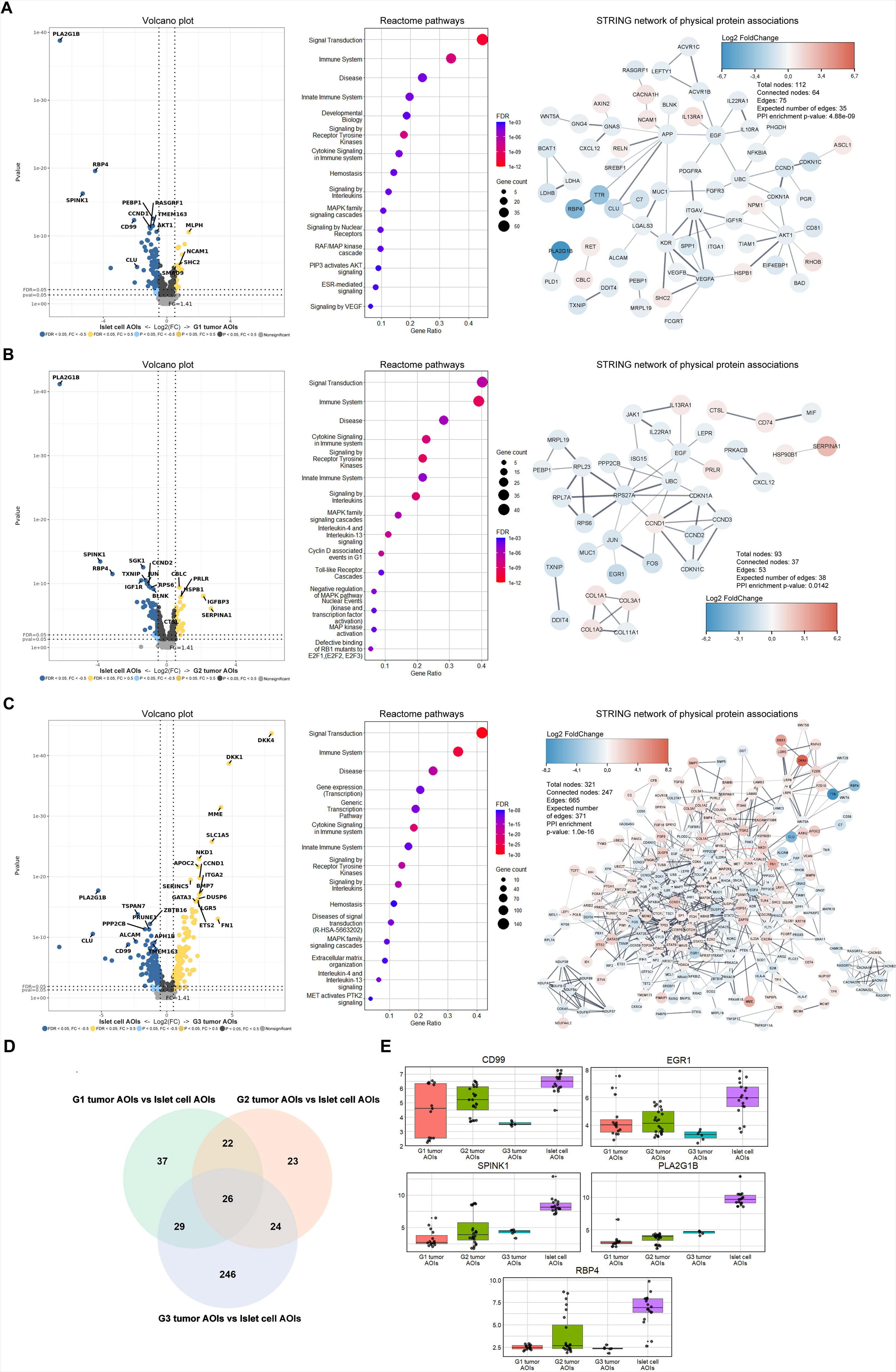
Comparison of tumor cell AOIs against pancreatic islet cell AOIs across all three tumor grades. **A - C** figures contain volcano plots depicting the Log_2_ fold changes and p-values of assessed genes, dot plots of the top 15 Reactome pathways according to FDR value which is sorted by gene count related to the term, and lastly, STRING network of physical protein associations depicting the physical interactions between proteins of DEGs. **A** - G1 tumors vs. islets; **B** - G2 tumors vs. islets; **C** - G3 tumors vs. islets. **D -** Venn diagram depicting the overlapping DEGs between three comparisons. **E -** Boxplots of Log_2_(Normalized counts + 1) of five consistently downregulated genes extracted from the list of top 10 downregulated genes across all three tumor grades.

In G2 tumors, the number of DEGs was slightly less than in G1 tumors (95 DEGs, 26 upregulated, and 69 downregulated). In enrichment analysis, these DEGs showed statistically significant associations with 168 Reactome pathways (Supplementary Table 9). Interestingly, the two downregulated DEGs genes related to the ubiquitin system (*UBC* and *RPS27A*) were present in more than half of 168 pathways, 116 and 131, respectively. Amongst the top 15 pathways (Figure 8 B), we found multiple pathways related to cell proliferation, differentiation, and survival (MAPK pathways, cyclin D-associated events in G1, defective binding of RB1 mutants to EF1). Amongst the 168 pathways, we did not detect pathways that are associated with VEGF signaling, which was prevalent in G1 tumors, as none of the DEGs related to VEGF signaling in G1 tumors were differentially expressed in G2 tumors. In a network of physical protein associations, 53 interactions between 37 proteins were found with a PPI enrichment p-value of 0.01.

In G3 tumor, the number of DEGs was the highest as 325 genes (152 upregulated, 174 downregulated) were dysregulated. Here enrichment analysis showed 320 statistically significant Reactome pathways (Supplementary Table 9). Amongst these were three pathways related to VEGF signaling, which constituted of four upregulated (*MAPK14*, *HSPB1*, *SHC2*, *FLT1*) and six downregulated (*KDR*, *PRKACA*, *PAK3*, *PRKACB*, *RHOA*, *VEGFA*) genes. Within the top 15 pathways (Figure 8 C) MAPK signaling pathway was once again present. Another proliferation-related pathway present in the top 15 pathway list is the disease of signal transduction by growth factor receptors and second messengers. This pathway constituted 19 upregulated and 15 downregulated genes, amongst which were *DKK1* and *DKK4,* which were the two most upregulated genes in G3 tumor. The network physical protein associations with G3 tumor DEGs showed a higher level of interactions than in G2 and G1 tumor networks, as 665 interactions between 247 proteins were found with a PPI enrichment p-value of 1 × 10^−16^.

### Comparison of PanNET cell AOIs between different tumor grades

Our final goal was to estimate the differences in gene expression in tumor cell AOIs across the three grades (Supplementary Table 10). As a result, by comparing G2 against G1 tumor AOIs, we identified 83 DEGs (43 upregulated, 40 downregulated) which showed significant associations with 41 Reactome pathways (Supplementary Table 11). The top 15 pathways (Figure 9 A), according to p-value, corresponded to immune system events, developmental biology (axon guidance, more specifically), cell proliferation, and estrogen-mediated signaling. In the STRING network of physical protein associations, 38 physical interactions between 37 proteins were identified at a PPI enrichment p-value of 0.001. Further inspecting all Reactome 41 terms amongst these four terms were related to VEGF signaling, which included two upregulated (KDR, VEGFA) and three downregulated (*NPR1*, *PRKACB*, *ROCK1*) genes in G2 vs. G1 tumors. In relation to cell proliferation, one upregulated (*CCND1*) and three downregulated (*CCND2*, *RBL2*, *CCND3*) genes in G2 vs. G1 tumors were identified. In string protein-protein interaction networks, these genes showed physical interactions with high confidence, and their associated Reactome terms were: Cyclin D associated events in G1, Generic Transcription Pathway, Mitotic G1 phase, and G1/S transition.

**Figure 9.**
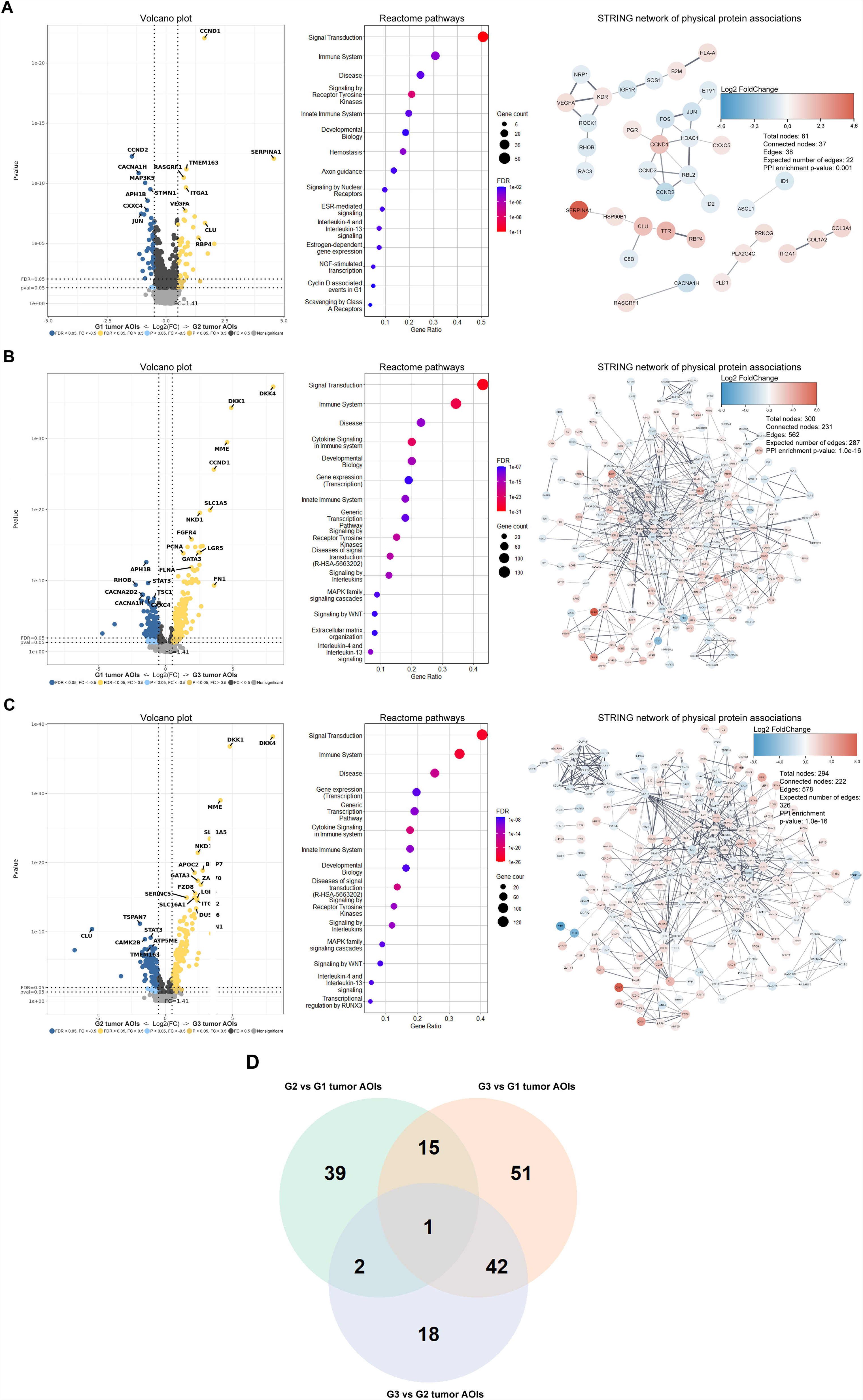
Comparison of tumor cell AOIs between different grades of the tumor. The figures contain volcano plots depicting the Log_2_ fold changes and p-values of assessed genes, dot plots of the top 15 Reactome pathways according to FDR value, and STRING network of physical protein associations depicting the physical interactions between proteins of DEGs. **A** - G2. vs G1; **B** - G3 vs. G1; **C** G3 vs. G2.

In both G3 vs. G1 and G3 vs. G2 comparisons, the amount of DEGs was substantially higher than in G2 vs. G1, with 307 (161 upregulated, 146 downregulated) and 294 (160 upregulated, 134 downregulated) respectively (Figure 9 D). In G3 vs. G1, the DEGs were associated with 261 Reactome pathways (Supplementary Table 11). Here amongst the top 15 Reactome terms (asides from immune system-related terms), MAPK family signaling cascades and signaling by WNT pathways were present (Figure 9 B). Both pathways included the two most upregulated genes in G3 tumor (*DDK1* and *DKK4*), which according to the Genecards database, antagonize canonical Wnt signaling by inhibiting LRP5/6 interaction with Wnt. A similar result was also observed for the G3 vs. G2 comparison, where both Wnt and MAPK pathways were present amongst the top 15 (Figure 9 C) of the 277 Reactome pathways (Supplementary Table 11). Here *DKK1* and *DKK4* genes were also the most upregulated. In G3 vs. G1 comparison, the proteins of DEGs showed 562 physical interactions between 231 proteins, and in G3 vs. G2 comparison, 578 interactions between 222 proteins. For both networks, the PPI enrichment p-value was 1 × 10^−16^.

## Discussion

In this study, using the DSP technology, we were able to segment the eight non-functional PanNET tissues and evaluate the gene expression of 1800 cancer-associated genes in three different compartments/AOIs α-SMA+ stroma, tumor adjacent to the α-SMA+ stroma and tumor non-adjacent to the α-SMA+ stroma. As a result, we were able to identify gene expression profiles characteristic of α-SMA+ stroma in PanNETs. Additionally, the DSP technology allowed us to identify and extract gene expression profiles specifically from pancreatic islets, which contain α, β, δ neuroendocrine cells. Since PanNETs are known to arise from either α or β cells of islets ^33^, the comparison of tumor AOIs against islet AOIs allowed us to more precisely investigate the changes in gene expression profiles present in PanNET cells.

The expression levels of α-SMA in the stromal compartment of the tumors have been linked to worse clinical outcomes and therapeutic resistance ^13,14,34^. One of the principal explanations for such observations is the fact that α-SMA is a marker for (CAFs) which arise from normal fibroblasts via TGF-β, osteopontin (OPN), interleukin-1β (IL-1β), PDGF, TNF-α, and FGF signals from the tumor ^35,36^. These signals, in turn, activate TGF-β, PI3K/AKT, Ras/MAPK signaling pathways, and NF-κB signaling pathways, leading to fibroblast transformation ^36,37^. Investigating these classical markers in our data, we did observe a statistically significant increase of *TNF* (TNF-α) in G1 tumors, *FGF14* in G2 tumors, and *FGF8*, *FGF18*, and *PDGFA* in G3 tumor. In addition, in our data regarding α-SMA+ stroma, we found that PDGF receptor beta (*PDGFR*) was overexpressed within α-SMA+ compartments in all three tumor grades. Additionally, the G3 tumor also had overexpression of PDGF receptor alpha (*PDGFRA*). The pathway enrichment analysis showed that *PDGFRB and PDGFRA* were involved in Reactome pathway R-HSA-2219528 (PI3K/AKT Signaling in Cancer), which was overrepresented in DEGs found in the α-SMA+ stroma of G2 and G3 tumors. The expression of PDGFRs has already been observed in stromal tissues of midgut NETs ^38^, and this data further adds layer of evidence that the PDGF/PDFGR axis may also exist in PanNETs. Another interesting observation is regarding the *NOTCH3* receptor gene in α-SMA+ stroma. There is rising evidence that cancer cells and fibroblasts interact directly through the cell surface ligand JAG1 and NOTCH receptors, and this, in turn, may also lead to fibroblast transformation into CAFs driving the process of CAF-mediated angiogenesis and extracellular matrix modification ^39,40^. A previous study in oral squamous cell carcinoma had shown that NOTCH3 is indeed overexpressed in CAFs, contributes to angiogenesis, and has been proposed as a therapeutic target ^40^. We observed that in our data, the *NOTCH3* receptor gene is overexpressed in α-SMA+ stroma compartments across all three tumor grades, and its ligand gene *JAG1* was found to be overexpressed in tumor compartments of G1 tumors. Considering the overexpression of NOTCH3 in α-SMA+ stroma compartments, we can assume the possibility that NOTCH may also be present in PanNET stroma.

Once fibroblasts are transformed and express α-SMA+, they can contribute to tumorigenesis in multiple ways. One of the main components that CAFs can regulate in TME is the extracellular matrix and immune infiltrates ^24,41^. One of the key genes that play a role in ECM modification is the stroma collagen gene ^42^. Stroma-derived collagens have been previously studied in terms of cancer progression, and in different combinations, they can either promote or inhibit tumorigenesis. For example, a study regarding PDAC showed that overexpression of type 1 collagen could promote tumorigenesis ^43^. Another *in-silico*-based study that analyzed multiple publicly available datasets identified collagen genes: *COL1A1*, *COL1A2*, *COL3A1*, and *COL5A1* as key CAF markers in gastric cancers ^44^. These same collagen genes were also upregulated in α-SMA+ stroma across all three tumor grades in our data. Besides the aforementioned collagen genes, we also observed overexpression of COL*5A2 and COL6A3* in G3 and G1 tumor/s. Another notable finding was gene *FN1* which encodes yet another stromal CAFs derived structural protein ^44^, and asides from being associated with invasiveness ^44^, some studies have shown that its expression positively correlates with immune infiltrates ^29,31,45^. In our data, we observe that the FN1 was upregulated across all three tumor grades, and its expression values were indeed associated with immune cell infiltrates of both myeloid and lymphoid lineage (Figure 5). Therefore, it would be valuable to see future studies on *FN1* and collagen genes in PanNETs as well to evaluate their role as prognostic biomarkers.

Aside from identifying the gene profiles and associated pathways within the α-SMA+ stroma of PanNETs, our other goal was to evaluate the impact of α-SMA+ stroma on tumor cells’ gene expression profiles. Since CAFs in the stroma and tumor cells communicate via cell-to-cell interactions and paracrine signals in both ways ^46^, we expected some differences in gene expression profiles of tumor cells immediately adjacent to α-SMA+ stroma and tumor cells non-adjacent to α-SMA+ stroma. This hypothesis has already been tested in an *in-vitro* study by *Wiechec et al.,* where the authors found 13 DEGs in 2D cultures of tumor cells co-cultured with CAFs in comparison to tumor cells cultured without CAFs and 81 DEGs in 3D cultures ^47^. While in our data, the differences are very marginal (Figure 7), we did identify two genes of interest: *MMP9* and *COL1A1*. These were the two most upregulated genes in G3 tumor compartments adjacent to α-SMA+ stromal cells; however, these results should be interpreted carefully as only the *MMP9* had an FDR value of < 0.05. Nevertheless, have been reported in a previous *in-vitro* study of a similar concept. Within the study, the authors found that *COL1A1* and *MMP9* were also amongst the most upregulated genes in tumor cells co-cultured with CAFs ^47^. Other studies have shown that these genes are involved in metastasis development ^45–47^, with MMP9 having a particularly vital role as it can degrade ECM and is one of the most widely studied metastatic markers ^48^.

Moving forward, we also looked at the gene expression profiles of the tumor itself. Since DSP technology allows us to investigate the expression within specific AOIs, we were able to compare the tumor compartments against pancreatic islets of Langerhans in adjacent normal tissue, which contain α and β cells, the deemed progenitors PanNETs ^49^. Currently, there are only a few studies that have used either the traditional bulk RNA-seq or microarray approach to investigate the PanNET tissues ^18,50,51^. Moreover, there are only two studies that have included the islets as reference samples when comparing the gene expression profiles ^52,53^. In our study, when we compared each grade separately, we first observed that five genes were consistently amongst the list of top 10 downregulated genes across all three tumor grades (Figure 6 E). These five genes have been linked to both pancreatic cancer and neuroendocrine tumors, particularly *CD99*, where the loss of it in PanNETs has previously been associated with a worse prognosis ^54–58^. Furthermore, the *RBP4* was also found to be downregulated in *Dilley et al.* study, where the authors also used islets as a reference ^53^. Though it is worth mentioning that the overall overlapping DEG count with this study was relatively low as different gene panels were used, and authors analyzed Multiple Endocrine Neoplasia type 1 PanNETs, whereas we analyzed sporadic tumors ^53^.

Analyzing the pathways overrepresented in the list of dysregulated genes across all three tumor grades, we identified a multitude of cancer-related pathways which have also been associated with neuroendocrine tumors ^59^. Of these, the notable pathways were R-HSA-2219528 (PI3K/AKT Signaling in Cancer), which was overrepresented in DEGs of G1 and G3 tumors, and R-HSA-157118 (Signaling by NOTCH) in G2 and G3 tumors. Another finding was the overrepresentation of MAPK signaling pathways, particularly in G2 and G3 tumors. MAPK signaling is implicated to play a role in tumor development as it is activated by growth factor receptors, and the resulting signaling cascades lead to cellular proliferation and survival ^60^. In neuroendocrine tumors, some attention has also been given to VEGF signaling pathways ^61^. In our data, the VEGF pathways were overrepresented in G1 and G3 tumors; however, the core gene *VEGFA* was downregulated in both G1 and G3 tumors. VEGF is known to play an essential role in neuroendocrine tumor development by stimulating angiogenesis ^17^. A study by *Zhang et al.* found that GEP-NETs tumor cells express VEGFA in higher levels compared to stroma ^62^ which was also the case in our study (Figure 5 A); however, our study shows that the expression levels of VEGF seem to be lower in PanNET cells than in islet cells. This result is not entirely unexpected as literature shows *VEGFA* is also critically important for islet cell development ^63^, and the comparison of VEGFA expression in islet cells and PanNET cells has not been studied yet. Lastly, signaling by WNT was also overrepresented in DEGs of G2 and G3 tumors. In G3 tumor regarding the WNT signaling pathways, we observed two genes that were two of the most upregulated DEGs (*DKK1* and *DKK4*). The Secreted Dickkopf (Dkk) proteins are known to be major elements in Wnt signaling, where they function as inhibitors of canonical Wnt signaling by blocking the interaction with LRP5/6 ^64^. The role of these proteins within tumorigenesis remains unknown, as recent literature seems to suggest that these Wnt antagonists can have both tumor-suppressive and oncogenic effects ^65^. Nevertheless, the overexpression of both DKK1 and DKK4 has been observed in pancreatic cancers and have been proposed as potential therapeutic targets ^66,67^.

Overall, the results of PanNET and islet cell comparison further support the evidence that the classical PI3K/AKT, NOTCH, MAPK, and VEGF signaling pathways may play a role in PanNET development, and it would be interesting to see further functional studies on these pathways in order to develop of novel treatment strategies.

After identifying the core gene expression profiles in stromal compartments and tumor compartments, we also carried out the comparison of both stroma and tumor across different tumor grades to evaluate the changes in gene expression of both compartments as the tumor advances. There is a limited amount of knowledge regarding the changes in gene expression within PanNETs as, according to our knowledge, there is only one bulk RNA-seq study by *Simbolo et al.* that did carry out the comparison between grades of resected PanNET tissues ^18^. In the study, it was discovered that the G3 vs. G1 tumor comparison had the highest amount of DEGs (2757), while G2 vs. G1 had the lowest (203). Based on these results, the authors concluded that G3 tumors are highly different from G1/G2 in regards to gene expression profiles as they identified a total of1104 overlapping DEGs between G3 vs. G1/G2 comparisons ^18^. Here we observe a similar result in our dataset as both α-SMA+ stroma and tumor compartment analysis had the lowest number of DEGs identified in the G2 vs. G1 comparison (58 in stroma, 83 in tumor) while the highest number of DEGs was in G3 vs. G1 comparison (109 in stroma, 307 in tumor). As for the G3 vs. G2, we observed 63 DEGs for stromal and 294 for tumor compartments which amounted to a total of 197 overlapping DEGs between G3 vs. G1/G2 comparisons, supporting the *Simbolo et al.* claim that G3 tumors have their own unique gene expression profile that is highly different from G1/G2 tumors.

The main limitation of this study is that we analyzed only one retrospective case of G3 tumor; this can be attributed to the fact that in a clinical setting, G3 tumors are the rarest form of PanNETs. The second limitation was encountered during the preparation steps as not all tumors were synaptophysin positive despite the fact that all tumors were histopathologically confirmed to express synaptophysin. As a result, for samples PanNET1, PanNET3, and PanNET4, the segmentation to acquire tumor cell AOIs was performed based on synaptophysin immunofluorescence signal, while for samples, the segmentation was performed based on morphology alone. Despite this, the cell deconvolution analysis showed a uniform distribution of tumor cell proportions except for the G3 tumor (PanNET8).

In summary, this study presents valuable information regarding PanNET biology, providing a cornerstone for future functional studies. Using the DSP approach, we were able to characterize PanNETs with a higher resolution as we analyzed α-SMA+ stromal cells and tumor cells separately. This allowed us to identify potential core genes in PanNET stroma (*COL1A1*, *COL1A2*, *COL3A1*, *COL5A1, COL5A2, COL6A3,* and *FN1*) and tumor (*RBP4*, *SPINK1*, *PLA2G1B*, *EGR1*, and *CD99*), outlay the altered molecular pathways and pinpoint the potential mechanisms of crosstalk between tumor and stroma (*PDGF/PDGFR* axis, *NOTCH3*, *MMP9*). We also show that as the PanNETs advance in grade, the gene expression profiles change not only in the tumor but in the stroma as well.

## Data Availability

All datasets containing raw/normalized counts and metadata information on each AOI analyzed in this study are described in the materials and methods section and are available in the supplementary material. Any additional datasets and information can be requested via the corresponding author/s.

## Acknowledgments

The authors acknowledge the Latvian Biomedical Research and Study Centre and the Genome Database of the Latvian Population and Instituto Ramón y Cajal de Investigación Sanitaria - IRYCIS for providing infrastructure, biological material, and data. During the study and manuscript preparation, Helvijs Niedra was supported by “SIA Mikrotīkls” patron; the donation was administrated by the University of Latvia Foundation.

## Ethics statement

The use of patients’ samples in this study was approved by the Central Medical Ethics Committee of Latvia (approval protocol No. 1.1–2/67) and Ramon y Cajal Ethical and Scientific Committees (protocol No. 196-19). The patients/participants provided written informed consent to participate in this study.

## Author Contributions

HN – carried out the digital spatial profiling analysis, bioinformatics analysis and manuscript preparation; RP and RS – performed additional data analyses and revisions; SV and JE – patient recruitment, clinical data collection and manuscript preparation; AG-B – patient recruitment and clinical data collection; AB and MGR – clinical data collection and tissue sample preparation; lastly, IR-C – assisted with tissue sample preparation, ROI selection during the digital spatial profiling analysis and manuscript revisions.

## Abbreviations

AOI: area of illumination
α-SMA: alpha smooth muscle actin
CAF: cancer associated fibroblast
CD99: Cluster of differentiation 99
COL1A1: type I collagen alpha chain 1
COL1A2: type I collagen alpha chain 2
COL3A1: type III collagen alpha chain 1
COL5A1: type V collagen alpha chain 1
COL5A2: type V collagen alpha chain 2
COL6A3: type VI collagen alpha chain 3
DEG: differentially expressed gene
DSP: digital spatial profiling
DKK: Dickkopf family proteins
FDR: false discovery rate
FGF: fibroblast growth factor
FN1: Fibronectin 1
G1/G2/G3: Grade 1/Grade 2/Grade 3
JAG1: Jagged1
MMP9: Matrix metallopeptidase 9
NET: neuroendocrine tumor
NOTCH3: Neurogenic locus notch homolog protein 3
TME: tumor microenvironment
PanNET: pancreatic neuroendocrine tumor
PDGF: platelet derived growth factor
PDGFR: platelet derived growth factor receptor
PPI: protein-protein interaction
RBP4: Retinol binding protein 4
RNA-seq: Ribonucleic acid sequencing
ROI: region of interest
TGF: tumor growth factor
VEGF: vascular endothelial growth factor

